# Optimisation strategies for the registration of Computed Tomography images of electropalatography

**DOI:** 10.1101/2020.04.02.022137

**Authors:** Agnieszka Sierhej, Jo Verhoeven, Naomi Rachel Miller, Constantino Carlos Reyes-Aldasoro

## Abstract

Electropalatography is a technique that employs a custom-made artificial palate to measure the contact established between the tongue and the hard palate. This technique is widely used in treatment of articulation disorders and studies of speech. In order to evaluate the accuracy of the electropalate, the device needs to be separated from the volume that usually contains electropalate worn on hard palate. This is done with the use of segmentation techniques. Prior to the segmentation, the registration of the two volumes, one containing the electroplate worn on hard palate, and one containing only hard palate, needs to be done. The registration is a technique of aligning multiple images by geometrical transform. Over the years, many methods for registration have been developed. The following paper describes the method of registration based on sensitivity analysis. Sensitivity analysis is a technique that evaluates the change in the number of pixels with different intensity with a shift of the volumes in different dimensions. Then based on the found optimal shift value, the shift in different dimension of the matrix is made. The technique successfully improves the alignments between two data sets, reducing the number of non-matching pixels. The sensitivity analysis-based registration should be useful in the future improvement of image processing tools that are crucial for the medical imaging.

## 1. Introduction

Electropalatography, also referred to as EPG, is a technique that allows to record the positioning and timing of the contact made between the tongue and the hard palate during the production of speech. EPG is a technique widely use in treatment of articulation disorder in adult population, and there are many studies to evaluate its usefulness in children population [1, 2]. A key aspect of EPG is the use of an artificial palate, which is a custom-made device used for collection of the data, which consists of a sequence of contacts (on/off) for a series of electrodes distributed over the palate [3]. Thus, it is crucial for the examination of articulation disorders that the electropalate is made with the highest accuracy and precision. It is possible to evaluate those by analysing the device and evaluating the placing of the electrodes. This can be done by scanning the electropalate with Computed Tomography and then analysing with medical image computing techniques. In order to separate the device from the stone cast of the palate, two sets of volumes can be used, one with the stone cast of the palate and one with the cast and the electropalate in position.

For almost half a century, the significant development in Medical Imaging has provided a series of techniques to observe the human body, and many other objects in ways not imagined before. Modern imaging modalities, such as magnetic resonance imaging (MRI), and computed tomography (CT), provide information about anatomy and physiology of the human body that can be use in treating many diseases. It is also useful to combine them with other modalities, e.g. positron emission tomography (PET). By registering image obtained from one modality to the images obtained from another, it is possible to acquire the deeper understanding of changes observed in physiology or anatomy of the body over the period of time[4].

This has posed new challenges to find new and better methods of analysing, comparing and combining the information acquired from different imaging modalities. Those needs have become the reason why the image registration is such a crucial technique in modern image processing [5]. Image registration is defined as the process in which multiple images are aligned through a geometrical transformation. This alignment, when optimally performed, allows the analysis of datasets and further steps to be performed. Image registration is also an essential technique when it comes to development of the segmentation techniques. An example of a wide group of segmentation techniques is a group of atlas-based segmentation where one volume is used as a reference volume to segment structure from a second volume[4]. It is necessary to register both volumes prior to the segmentation in order to make the segmentation more accurate. Over the years, a large number of techniques were developed and most of them rely on intensity-based approach. One of the most important question that can be asked while considering the registration is how to quantify the quality of the registration.

This paper presents a methodology of image analysis steps, which include semi-automatic registration and sensitivity analysis for the segmentation of electropalates.

## 2. Materials and Methods

This section describes a methodology in which computed tomography (CT) images of stone casts of human palates with and without an electropalate, (artificial palate used to examine the contact made between tongue and palate [6]) are registered in order to segment the electropalate from the palate.

### 2.1. Materials

A series of electropalates [Figure 1] were scanned with a CT Scanner. The images were acquired with Soredex Scanora3D Cone Beam CT scanner. Four hundred and fifty-one DICOM images with the resolution 451×451 [pixels] and slice thickness of 0.133 [mm] were acquired at the power of 85 [kVp]. Data Set 1 contains images where the electroplate is worn, and Data Set 2 contains images of the palate of the alone.

**Fig. 1.**
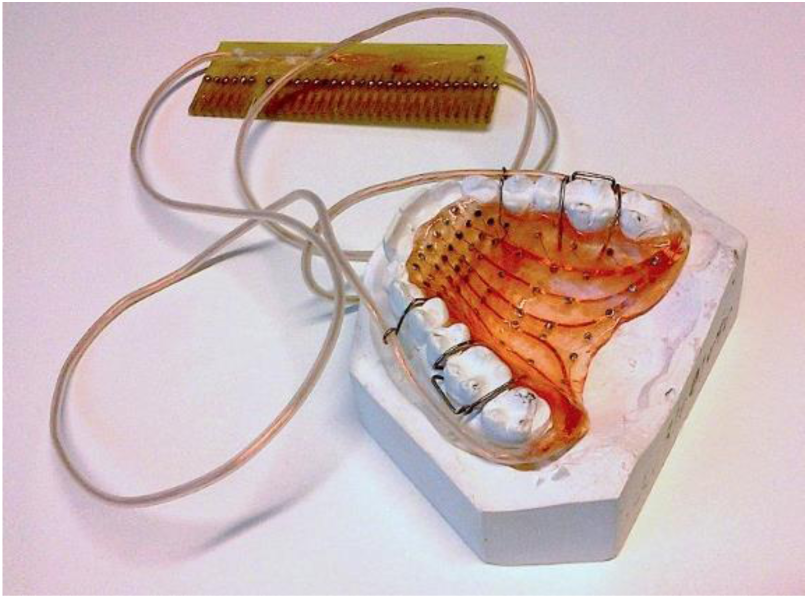
An electropalate mounted on a stone cast of a human palate. The figure shows the stone cast in white, with the palate and its electrodes mounted on the cast. Wires connect the electrodes to an external interface which is connected to a computer to read the signals from the electropalate.

### 2.2. Methods

In order to separate the device (electropalate) from the images, both data sets need to be perfectly aligned and then subtracted from each other. Initial misalignment of both data sets is shown in Figure 3, where white and black areas represent the misaligned areas and grey area presents perfect alignment of the pixels with the same intensity. It needs to be emphasised that some of the white areas represent the electrodes of the device, however those are not affected by the subtraction of both data sets.

**Fig. 2.**
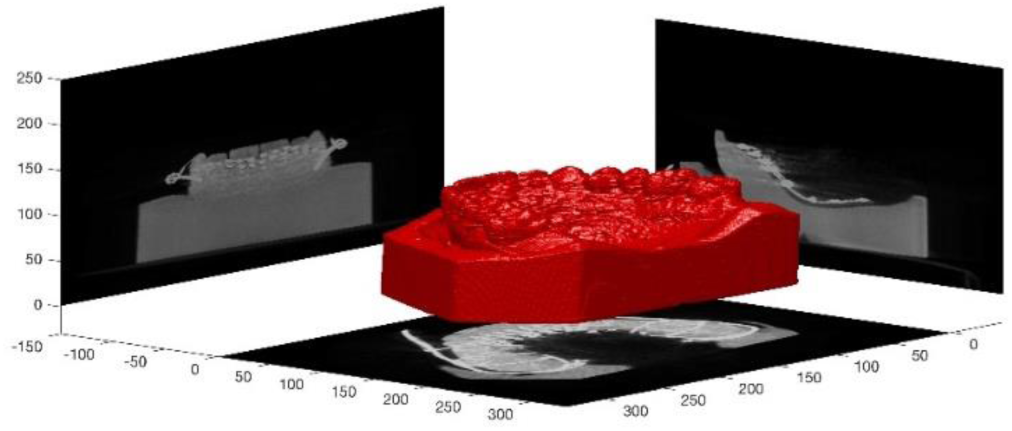
An illustration of the data acquired with Computed Tomography. Maximum intensity projections of each orientation are shown with a 3D rendering of the electropalate and the stone cast.

**Fig. 3.**
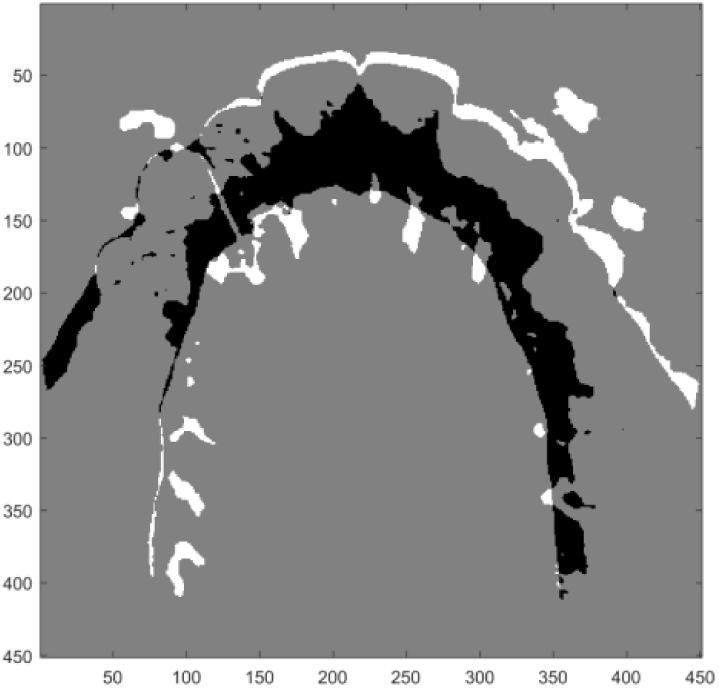
Illustration of the initial misalignment of the Data Set 1 and Data Set 2. Black and white areas denote the pixels with non-zero value of intensity after the subtraction of both volumes, whereas the grey area denotes the pixels with zero intensity, which were perfectly aligned between both data sets. Some of the white areas denotes the electrodes.

Using the sensitivity analysis, which measures the difference in number of pixels after subtracting matrix of Data Set 2 from matrix of Data Set 1, it is possible to examine which shift value have the best impact on quality of the registration, using quantification methods.

### 2.3. Levels, rows and columns (z-, x- and y-dimension) shift of 3D matrix

Data Set 1 is set will be considered as a fixed set, while Data Set 2 is set as a moving set, which will be displaced along three dimensions: levels, rows and columns [Figure 4]. The algorithm analyses two shifts: forward (positive values) and backwards (negative values). Using the for loop, the chosen slice of the matrix from Data Set 2 is shifted and the shift is compared with fixed slice form Data Set 1 by subtracting both slices from each other.

**Fig. 4.**
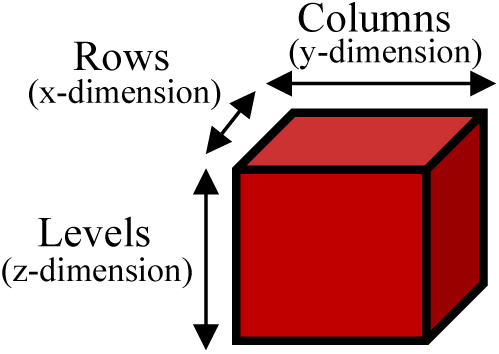
Description of levels, rows and columns (z-, x- and y-dimension, respectively) of the 3D matrix used in this work.

Sensitivity analysis is based on the following steps. The sum of absolute value of all pixels is calculated. If the values (intensities of the pixels) from both data sets match perfectly, the difference is zero. It is expected that the lower the sum, the greater quality of alignment is, as more pixels with the same intensity in both data sets aligns. The values of difference in intensity for each shift value are recorded and stored in a separate vector. Then, the values are analysed to find the minimum, which in this case, is the optimal value of the shift. The algorithm automatically selects the lowest value of difference in intensity and the corresponding shift value, which then, becomes the optimal shift value. If the optimal shift value is equal to zero, the new matrix becomes the previous matrix. If the optimal shift value is different from zero, the appropriate shifting algorithm is automatically applied to and the new matrix is computed. Same procedure is applied to rows (x-dimension) and columns (y-dimension).

Sensitivity analysis graphs [Figure 5] shows that in the case of those data sets, the misalignment was present in levels [Figure 5a] and in columns [Figure 5b], but not in rows [Figure 5c].

**Fig. 5.**
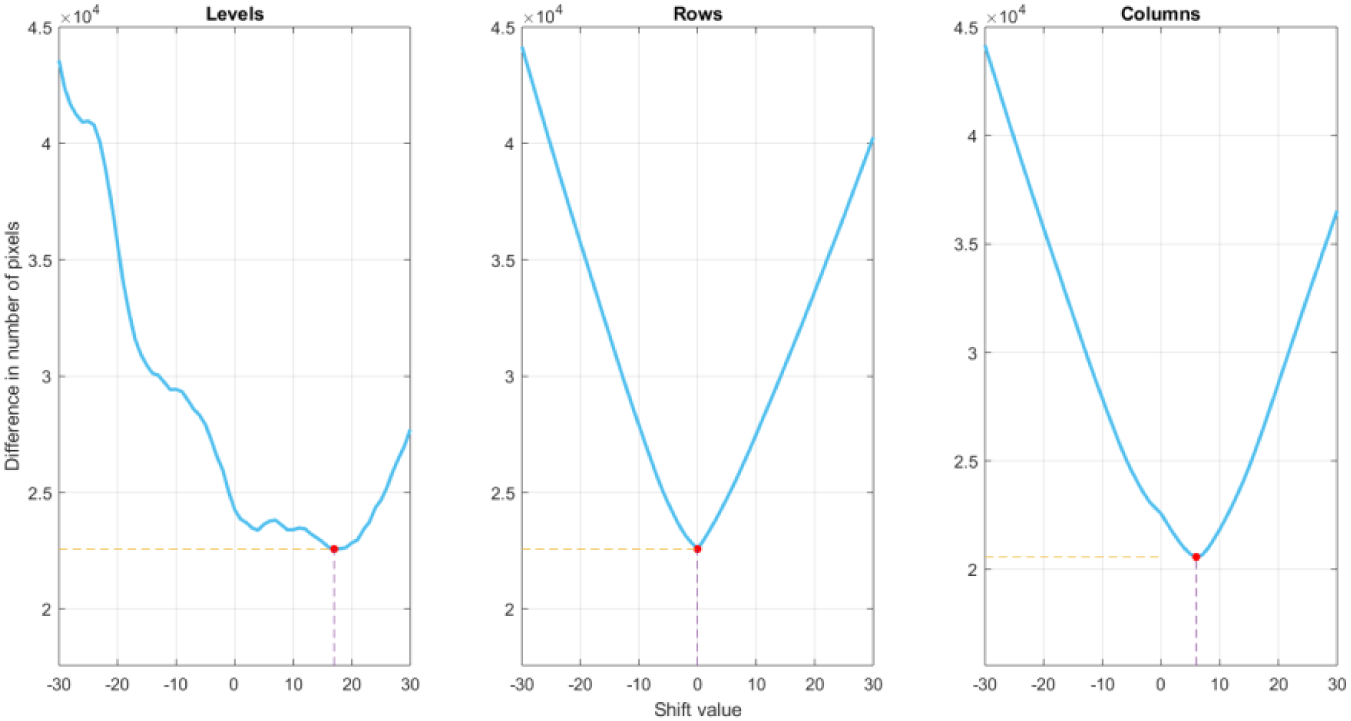
Sensitivity analysis for (a) levels, (b) rows, and (c) columns. Shift value denotes the number of levels, rows, or columns that have been shifted, either forward (positive values) or backwards (negative values). The difference in number of pixels shows how many pixels with non-zero intensity is left after the subtraction of both volumes. Red dot shows the position of optimal shift value in columns

### 2.4. Angular shift

The angular shift algorithm is slightly different form algorithm that is utilized in levels, rows and columns shift. The reason behind existing, minor changes in the algorithm is the limits of the MATLAB command ‘imrotate’ that was used in this section. Command ‘imrotate’ is used to rotate matrixes, by a chosen angle, around the centre of rotation which is set to me the centre of the entire matrix. However, in the case of those data sets, the centre of rotation has to be placed in the landmark which is the axial slice of one of the teeth. The centre of rotation, however, must be specified by inspection, which is done manually rather than automatically. The specific coordinates must be chosen so the algorithm could expand the matrix to shift the centre of rotation.

The original centre of rotation can be found in coordinates (*x*_*original centre of rotation*_, *y*_*original centre of rotation*_) calculated by the following equations:

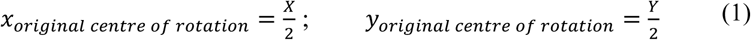

Where: X - size of the original matrix in x-dimension, Y - size of the original matrix in y-dimension.

It is assumed that in the case of those data sets x _new centre_ is always greater than 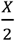, because the obtained landmark (one of the teeth) is placed always in the ‘upper half’ of the x-dimension. Therefore, x _added_, which denotes number of rows containing ‘0’ that needs to be added to the matrix in order to shift the centre of rotation, will be always present on one side of the matrix, and never on the other one. However, to find the y _added_, which denotes number of columns containing ‘0’ that needs to be added to the matrix in order to shift the centre of rotation, the two cases needs to be considered [Figure 6].

**Fig. 6.**
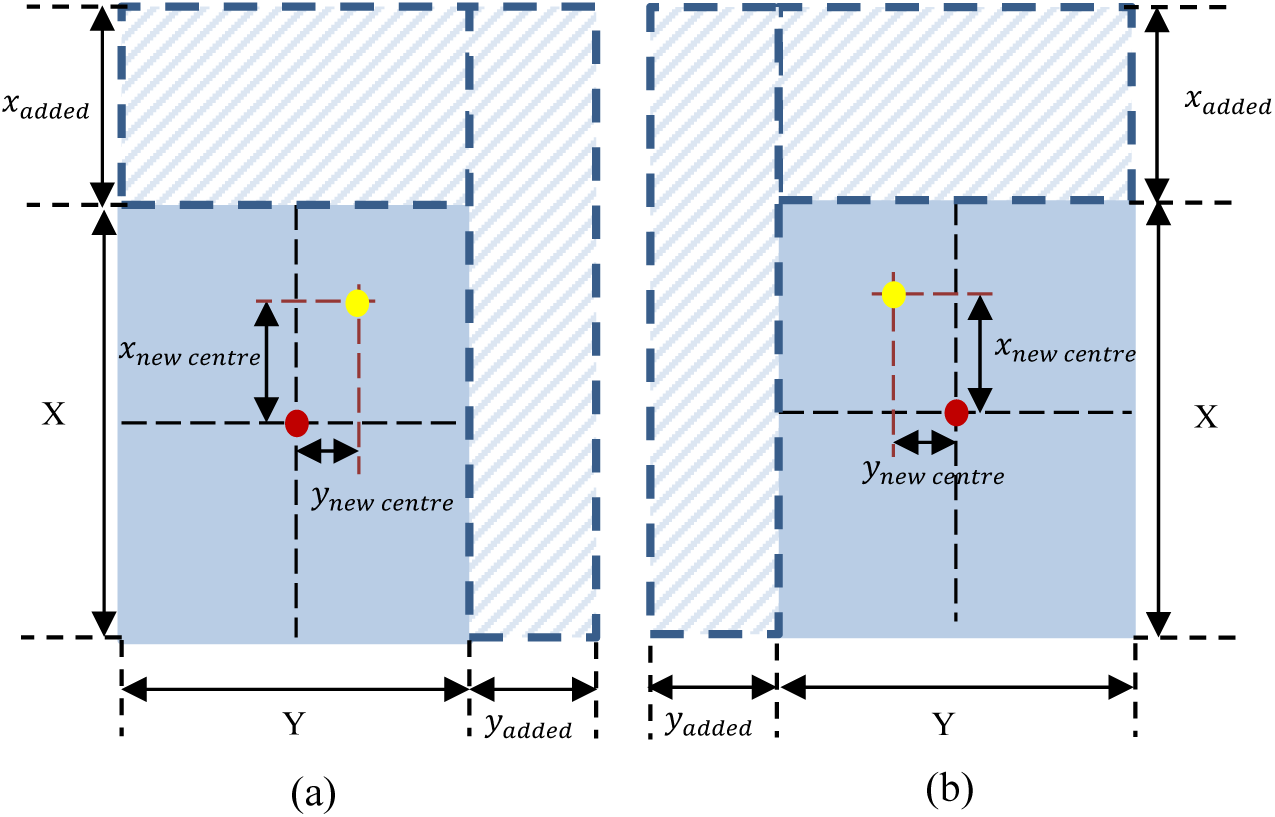
Representation of the (a) first and (b) second case of shifting the centre of rotation. Red dot represents original centre of rotation, whereas yellow dot represents new centre of rotation. Blue area represents the original matrix, and the area with diagonal lines represents the rows and columns added to the matrix.

The first case (a) assumes that y _new centre_ is greater than 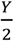. In that case, the following equations are used to find the coordinates of the new centre of rotation:

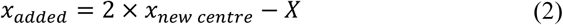

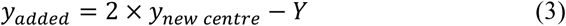

The second case (b) assumes that y _new centre_ is smaller than 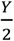. In that case, the following equations are used to find the coordinates of the new centre of rotation:

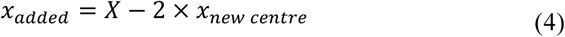

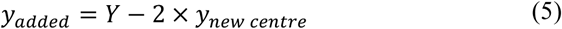

It is worth to note that only x-dimension and y-dimension are taken under consideration in finding the coordinates of new centre of rotation. It is due to the fact that the command ‘imrotate’ rotates all slices of the 3D matrix, according to those coordinates. Therefore, z-dimension can be omitted in those considerations.

After finding the values of x _added_ and y _added_, a new matrix is computed, which contains values form the old matrix, but has greater dimensions than an old matrix (greater by the values of x _added_ and y _added_). Those additional rows and columns are filled with ‘0’, with the purpose that they will be removed after the shifting. For the testing purposes, Data Set 1 matrix is rearranged as well in the same manner.

Following the rearrangement of both matrices, the two consecutive loops are performed to search for the best alignment for the angular shift. The values from −10° to +10° are investigated. The optimal value of angular shift is found and the search for it is visualised in sensitivity analysis for angles [Figure 7].

**Fig. 7.**
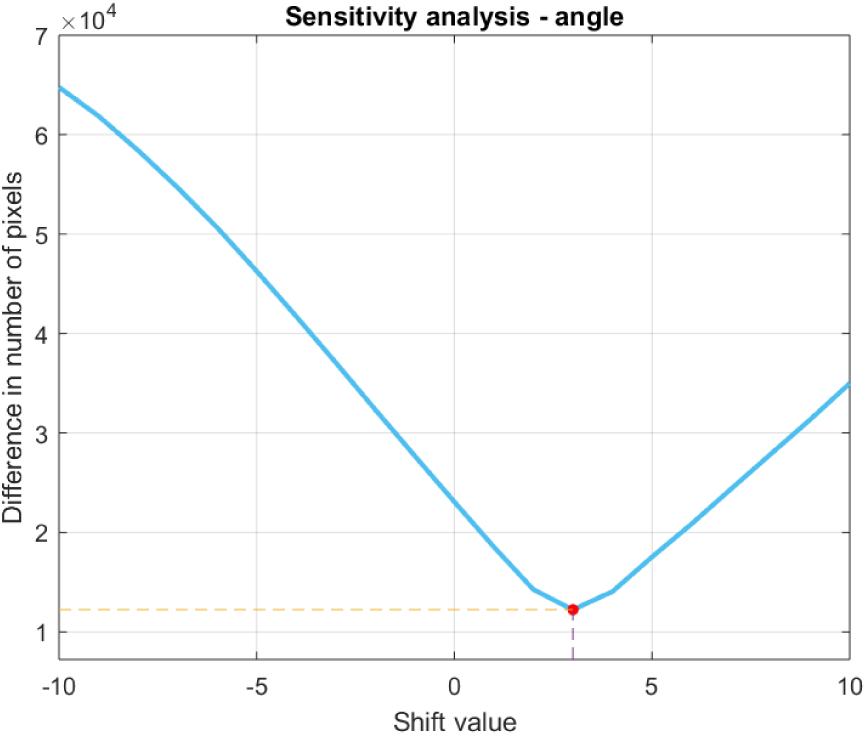
Sensitivity analysis for angles. Shift value denotes the rotation with the specified angle, either forward (positive values) or backwards(negative values). The difference in number of pixels shows how many pixels with non-zero intensity is left after the subtraction of both volumes. Red dot shows the position of optimal shift value in angle.

The new matrix with better alignment is computed, and then the additional rows and columns are removed, thus the new matrix has the same dimensions as the previous matrix. The significant improvement in the alignment can be observed by comparing the black and white areas in Figure 8a and Figure 8b. It is worth to note that the white areas which represents the electrodes (and wires), were not affected by the registration.

**Fig. 8.**
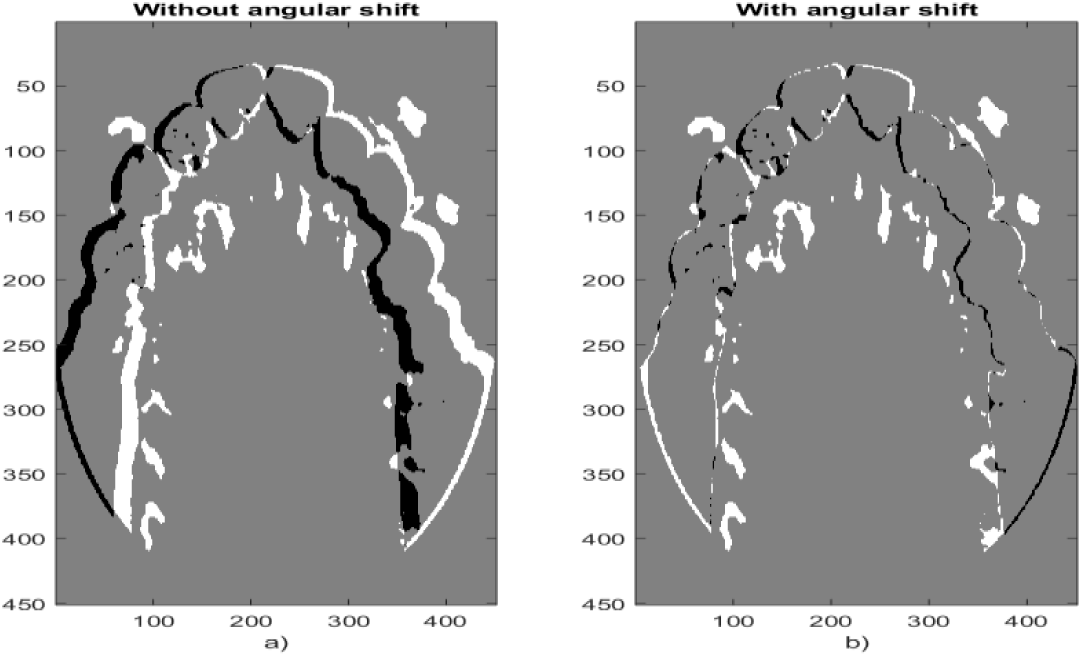
Difference in pixels with (a) or without (b) shift in angles. The black or white area shows there is a difference in pixels value. Grey area denotes that the difference in pixels is zero.

### 2.5. Second analysis for rows and columns dimensions

Analysing the alignment after angular shift [Figure 8b], the following assumption can be made: the further improvement of alignment is still possible. Therefore, the second analysis of rows and columns is performed. The following figures confirms the assumption that has been made. Although there is no need for further shift in columns dimension, the shift in rows dimension should be made in order to accomplish better alignment.

Figure 10 presents the effect of complete registration on alignment of both data sets. It is clear that without registration, there is a great misalignment between both data sets, which can be seen as large white and black areas in Figure 10a. Those areas represent difference in intensity of the pixels of both data sets, therefore, the smaller they are, the better is the alignment. The improvement registration allows to segment the device from the data sets [Figure 11].

**Fig. 9.**
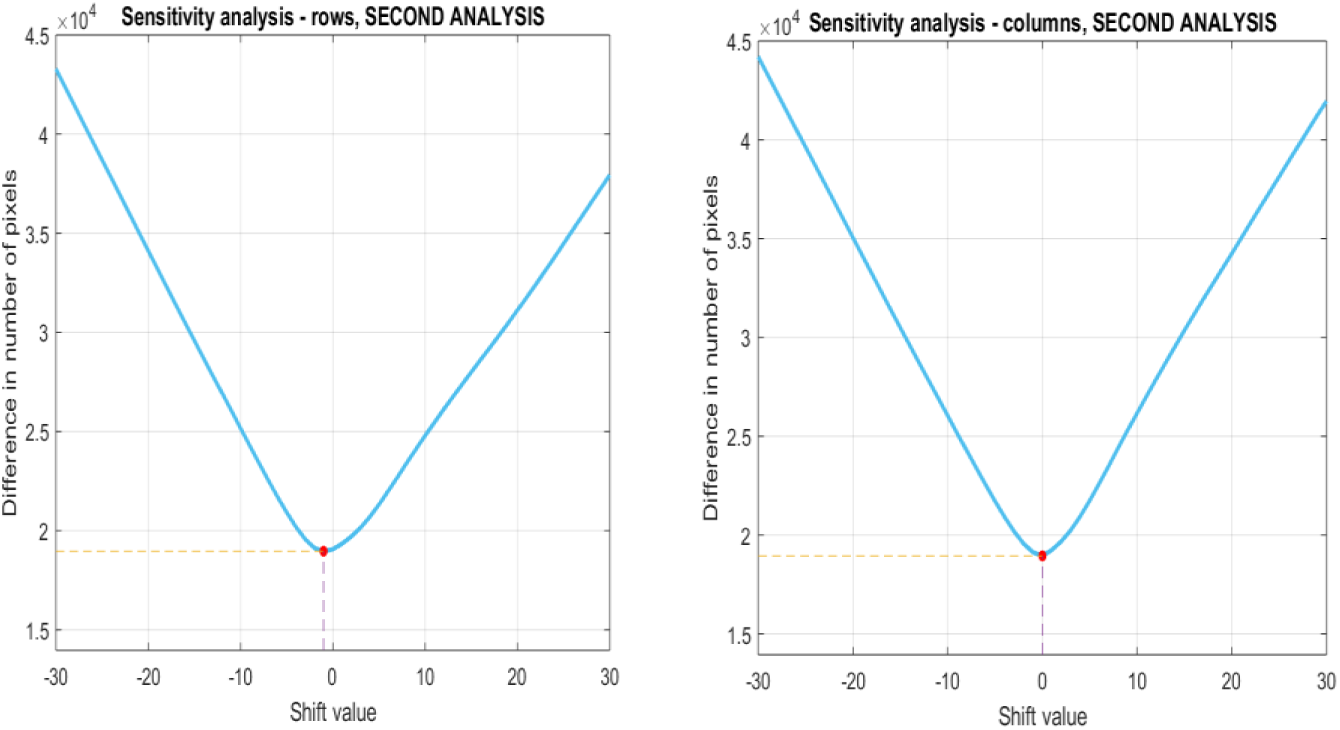
Second sensitivity analysis for (a) rows and (b) columns. Red dot shows the position of optimal shift value in columns

**Fig. 10.**
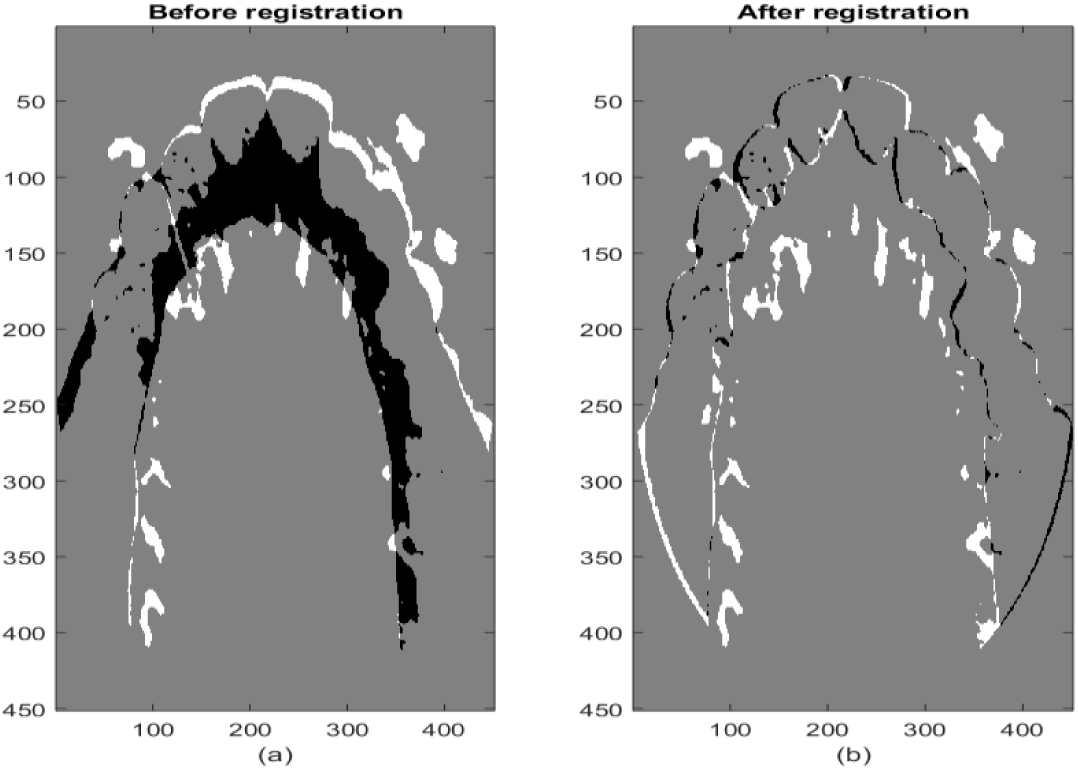
Comparison of alignment between data sets (a) before registration and (b) after registration was completed.

**Fig. 11.**
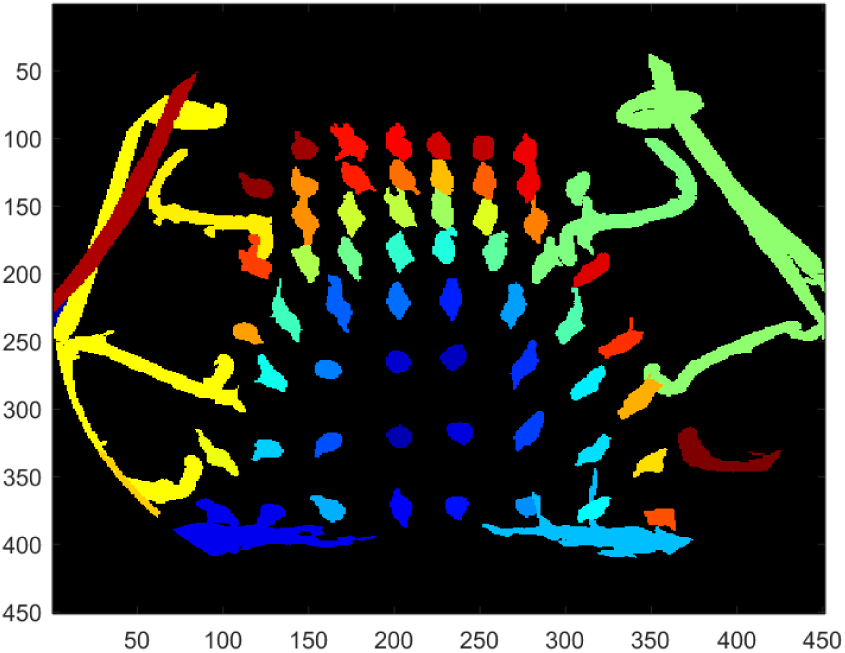
Final segmentation of the electropalatogram. Wires and electrodes are labelled with different colours for visualisation purposes. It should be noticed that there are some minor artifacts, which may be removed with further post-processing.

## 3. Discussion

The registration process by shifting in levels, rows, columns, and angularly, presented above successfully improved the alignment between two data sets, without negatively affecting the electrodes (and wires). The white and black areas, which represents the pixels with non-matching intensity, are largely reduced.

It has to be emphasised that the initial misalignment is often hard to observer when analysing the two volumes separately. It is more convenient and accurate to analyse them combined together, to observe the true level of misalignment. As it is presented in the Figure 3, the initial alignment exists but may not be obvious without combining and comparing two data sets. Thus, it is important to evaluate the alignment in any situation, while dealing with data sets which needs to be segmented or combined to be examined together.

In contrast to the Figure 5b and 5c, Figure 5a presents the curve with unregular shape. The analysis emphasises that choosing the shift by visual inspection can be highly misleading as a wide range of shift values gives relatively good improvement of alignment. However, in order, to choose the best alignment, the numerical evaluation needs to be performed. Figure 5 shows that in the case of those data sets the most significant misalignment is observed in levels dimension, however, in the case of any different data sets, this may vary. Analysing Figure 5b and 5c, it can be stated that the scans of those data sets were taken with the high precision according to some landmarks as the reference point, where two volumes were almost perfectly aligned. However, this may vary for different data sets. Current method analyses the shift only in one dimension, starting from levels dimension. Future work on the method needs to include the evaluation process of the order in which dimensions should be analysed.

Angular shift process requires of finding the new centre of rotation. It was highly automated by this method; however, the present method still requires choosing manually the new coordinates. The method has a crucial disadvantage as it is based on the subjective assessment by the observer, which may vary from one person’s point of view to another. The studies performed in the last decade showed that there is a large issue with reliable identification of the landmarks. Nevertheless, some improvements in this field were achieved recently and further work can be done in the future [7, 8]. The issue may be solved by deep-learning regression networks methods, which are currently developed, and recent research concludes that they may have great potential in the field of medical imaging [9]. Moreover, the method assumes that the new centre of rotation is placed in the “upper half” of the matrix. Future developments should investigate the cases where the new centre of rotation can be also found in the “lover half” of the matrix.

Careful observation of Figure 7 may show that even 1degree rotation highly affects the alignment, making the angular shift very sensitive. From Figure 8, it can be observed that the angular shift is highly crucial for the improvement of alignment. However, as it was stated previously, the further improvement can be made. Observation that can be made while analysing Figure 9 clearly proofs that. It also proofs that the future improvement for the code has to be made so it evaluates the alignment and find the optimal value shift in all dimension for multiple dimensions and not just one dimension at time [10]. The biggest disadvantage of that could be the time of running the code, but this could be solved by improvement made in both the hardware and software.

Figure 10 illustrates that the registration based on sensitivity analysis is successful. The final segmentation of wires and electrodes is shown in Figure 11, which confirms the success of the methodology. It should be highlighted that past the alignment, the segmentation is far simpler than an earlier segmentation methodology presented by Verhoeven [6]. Nevertheless, further improvements to this method could be made. Future action may include multi-dimensional sensitivity analysis as oppose to the one-at-the-time dimension sensitivity analysis. All the work presented in this paper can be used to improve many image processing techniques, which are nowadays crucial in medical imaging.

